# Metamorphosis imposes variable constraints on genome expansion through effects on development

**DOI:** 10.1101/2021.05.05.442795

**Authors:** Rachel L Mueller, Clayton E Cressler, Rachel S Schwarz, Rebecca A Chong, Marguerite A Butler

## Abstract

Genome size varies ~ 100,000-fold across eukaryotes and has long been hypothesized to be influenced by metamorphosis in animals. Transposable element accumulation has been identified as a major driver of increase, but the nature of constraints limiting the size of genomes has remained unclear, even as traits such as cell size and rate of development co-vary strongly with genome size. Salamanders, which possess diverse metamorphic and non-metamorphic life histories, have the largest vertebrate genomes — 3 to 40 times that of humans — as well as the largest range of variation in genome size. We tested 13 biologically-inspired hypotheses exploring how the form of metamorphosis imposes varying constraints on genome expansion in a broadly representative phylogeny containing 118 species of salamanders. We show that metamorphosis during which animals undergo the most extensive and synchronous remodeling imposes the most severe constraint against genome expansion, with the severity of constraint decreasing with reduced extent and synchronicity of remodeling. More generally, our work demonstrates the potential for broader interpretation of phylogenetic comparative analysis in exploring the balance of multiple evolutionary pressures shaping phenotypic evolution.

## Introduction

Across the tree of life, few traits exhibit the tremendous scale of variation shown by genome size, which encompasses a ~100,000-fold range across eukaryotes alone (Gregory 2022). Decades of research have explored the question of whether this variation has a cohesive evolutionary explanation, revealing two traits that consistently covary with large genome size: cell division rate slows down and cell size increases (Gregory 2001, 2005). However, whether genome size evolves by adaptation or constraint, and what drives these processes, have been challenging questions to answer, in part because the organismal features involved have been unclear. For example, genome size co-varies in a context-dependent manner with metabolic rate, showing an association within some vertebrate clades, but not others, and shows no correlation across vertebrates as a whole (Licht and Lowcock 1991; Gregory 2002a; Smith, et al. 2013; Wright, et al. 2014; Kapusta, et al. 2017; Uyeda, et al. 2017; Gardner, et al. 2020). Genome size has also been associated with developmental rate or complexity (Gregory 2002b), temperature (Hessen, et al. 2013), invasiveness (Pandit, et al. 2014), and speciation and extinction rates (Vinogradov 2004; Jeffery, et al. 2016). As these associations vary in their generality, the explanation may not lie in a single overarching factor. Rather, genome size variation may reflect the balance of multiple evolutionary pressures. Thus, to disentangle the forces affecting genome size requires close consideration of the interaction of ecology and organismal biology (Gregory 2004; Roddy, et al. 2019).

The association between genome size and metamorphosis has been repeatedly noted across ectotherms (salamanders: Larson 1984; Wake and Marks 1993; Gregory 2002b; Sessions 2008; fish and insects: Gregory 2002b). It has long been proposed that natural selection acts to shorten the duration of metamorphosis (Szarski 1957) to limit exposure to potentially lethal stresses during transformation, termed “metamorphic vulnerabilities” (Gregory 2002b; Lowe, et al. 2021). Salamanders have received special focus because they have the largest genomes among vertebrates (9 Gb – 120 Gb per haploid genome; the vast majority are diploid), as well as exceptional diversity in life history across the 773 extant species (Decena-Segarra, et al. 2020; AmphibiaWeb 2022; Gregory 2022). Metamorphosis has been lost, modified, and regained throughout the clade’s evolutionary history leading to metamorphosers, paedomorphs, and direct developers among extant salamanders. These three life history types vary widely in the degree and duration of transformation, as well as many other organismal features that have important consequences for natural selection. Metamorphosers undergo a morphological transformation from an aquatic larval to terrestrial adult form, paedomorphs retain the aquatic larval morphology throughout life, and direct developers hatch from terrestrial eggs as miniature versions of the adults (Rose 1996; Rose 1999; Chippindale, et al. 2004; Mueller, et al. 2004; Wiens, et al. 2005; Bonett, et al. 2014).

The observation that metamorphosis is linked to smaller genome size (Wake and Marks 1993; Gregory 2002b; Sessions 2008; Bonett, et al. 2020), and larger genome size is linked to an overall slow-down of developmental rates, led to the hypothesis of time-limited metamorphosis as a constraint on genome expansion (Gregory 2002b). An evolutionary constraint is broadly defined as a limit to phenotypic variation (Arnold 1992) that can arise from non-adaptive sources or from indirect selection on a correlated trait (e.g., Savell, et al. 2016). For example, because the rate of development is an emergent property of complex cellular processes that are slowed down by large genome/cell sizes (Horner and Macgregor 1983; Jockusch 1997), selection to shorten metamorphosis should limit the sizes that genomes can attain, resulting in a constraint on genome size. There has been some ecological confirmation of the time-limited metamorphosis hypothesis in salamanders, with species in ephemeral habitats possessing smaller genomes (Lertzman-Lepofsky, et al. 2019). However, large-scale phylogenetic comparative analysis has both rejected (Liedtke, et al. 2018) and confirmed the classical association between genome size and life history (Bonett, et al. 2020). A key difference in the latter was partitioning life history diversity to better reflect how developmental complexity influences genome size. Combining lineages with the ability to metamorphose at all even if only occasionally, as seen in facultative paedomorphs — and separating them from lineages that are obligate paedomorphs was key to finding any association between genome size and life history (Bonett, et al. 2020).

Although this was a big step forward, there was no clear best model; a model that separated direct developers from paedomorphs and a model that grouped them as non-metamorphosers explained the data nearly equally well. This result is surprising if developmental remodeling is a primary factor influencing genome size because direct developers undergo a substantial amount of metamorphic remodeling inside the egg. We hypothesize that this result may be explained by the models not yet considering natural diversity across salamander lineages in the synchronicity and extent of developmental remodeling during metamorphosis, as well as the associated vulnerabilities. Thus, the relevant features of metamorphosing organisms, and the evolutionary pressures they exert on genome size, likely remain incompletely understood.

In this study, we explore how metamorphosis shapes genome size, expanding the consideration of life history diversity to include natural variation in developmental complexity during metamorphosis and associated metamorphic vulnerabilities. We inform stochastic models of genome size evolution with life history data and the molecular mechanisms of genome expansion. We detail how OU model-based comparative methods can be used to explore adaptation and constraint in genome size and, by extension, in other traits that also may not be shaped exclusively by adaptive evolution. Whereas OU models have been used widely to study adaptive significance (Hansen 1997; Butler and King 2004), some traits such as extremely large genome size have no known fitness benefit, which is at odds with an interpretation of adaptive evolution. Indeed, these general models can be interpreted in multiple ways, and rather than evolutionary “optima,” the phenotypic locations predicted by the models may be better envisioned as “equilibria” to reflect a balance of forces such as upwardly-biased mutation pressure opposed by a constraint set by other aspects of organismal biology. The sigma term, describing the intensity of random fluctuations of the evolutionary process, has received much less attention, but in combination with the other parameters may be helpful in diagnosing release from constraint.

## Methods

### Genome Size and Life History Regimes

#### Stochastic Genome Expansion

In vertebrates, genome size is strongly shaped by transposable elements (TEs), sequences that replicate and spread throughout host genomes, increasing genome size (Sotero-Caio, et al. 2017; Shao, et al. 2019). TEs can be deleted by errors in replication, recombination, and DNA repair (Michael 2014; Vu, et al. 2017). TE activity can be nearly neutral, largely missing functional genomic regions and therefore resulting in negligible fitness consequences (Arkhipova 2018). Salamanders are particularly prone to stochastic increases in genome size through TE accumulation (Sun, et al. 2012; Keinath, et al. 2015; Nowoshilow, et al. 2018) because they possess low TE deletion rates and incomplete TE silencing (Sun and Mueller 2014; Frahry, et al. 2015; Madison-Villar, et al. 2016). Against this clade-wide background of biased stochastic genome expansion, we hypothesize that different life history regimes exert varying levels of constraint on genome size mediated through exposure to metamorphic vulnerabilities as explained below.

#### Metamorphic Vulnerabilities

Metamorphosis is a radical morphological transformation often coordinated in a short window of time, and thus subject to multiple risks. During metamorphosis, animals undergo rapid cell division, differentiation, migration, and apoptosis (Alberch 1989), as they remodel their skin and glands, blood, gut, teeth, and musculoskeletal, excretory, and immune systems (Rose 1999). Accordingly, metamorphosing animals can experience three classes of vulnerabilities: 1) <u>Performance and physiological handicap</u>. Animals can suffer reductions in performance in escaping predation or accessing food and shelter; or reductions in physiological tolerance to environmental swings during transformation. 2) <u>Energetic limitations</u> imposed when resorbing or remodeling larval structures and forming adult structures (Wassersug and Sperry 1977; Orlofske and Hopkins 2009; Enriquez-Urzelai, et al. 2019; Lowe, et al. 2019; Lowe, et al. 2021). 3) <u>Random developmental errors</u> resulting from developmental system perturbation, which can become more likely with both increased developmental complexity and reduced cell numbers when remodeling occurs at earlier embryonic stages (Hanken 1984; Gregory 2002b; Rose 2003).

In addition to the ancestral form of metamorphosis, salamanders have evolved a derived mode that differs markedly in timing and extent of remodeling (Alberch, et al. 1985; Rose 1999, 2003; Beachy, et al. 2017) In contrast to previous studies, we therefore differentiate four life history regimes for the evolution of genome size: Paedomorphosis, Direct Development, and two forms of metamorphosis called Gradual Progressive Metamorphosis and Abrupt Synchronous Metamorphosis. Because they account for < 2% of salamander species, we did not include viviparous and ovoviviparous life histories.

#### Paedomorphosis

Individuals do not undergo metamorphosis, instead reaching sexual maturity retaining largely larval traits without any radical developmental remodeling (Gould 1977; Rose 1996). Paedomorphosis has evolved at least eight times within salamanders (Rose 1999). Paedomorphs experience no metamorphic vulnerabilities and are thus predicted to be free from any related constraints on genome expansion.

#### Direct development

The larval growth stage is eliminated, and metamorphosis is integrated with embryogenesis into a single sequence of developmental events that takes place much earlier in ontogeny, inside the egg (Alberch 1989; Rose 2014). Thus, metamorphic remodeling occurs, but the events involve small amounts of tissue and few cells. Direct development has likely evolved at least twice within salamanders (Bonett, et al. 2014). Direct developers experience the metamorphic vulnerabilities of energetic limitation and developmental error due to decreased cell numbers. Thus, direct developers are predicted to have intermediate levels of constraint on genome size associated with metamorphosis.

#### Gradual, progressive metamorphosis

This type of metamorphosis is likely ancestral for salamanders (Rose 1999), and involves remodeling events happening sequentially and over relatively long timeframes in a free-living, aquatic organism in preparation for transition to a terrestrial habitat (Rose 1996). In contrast to frogs, these salamander larvae are able to feed throughout the process, as well as use larval energy stores (Semlitsch, et al. 1988; Deban and Marks 2002), alleviating energetic limitations. Furthermore, the timing of metamorphic onset is flexible and can be delayed when larval food resources are limited (Beachy, et al. 2017). Gradual metamorphosers experience the metamorphic vulnerabilities of decreased performance and physiological tolerance and are thus predicted to have intermediate levels of constraint on genome size associated with metamorphosis.

#### Abrupt, synchronous metamorphosis

This form of metamorphosis differs markedly from gradual, progressive metamorphosis in that remodeling events are more radical, occur simultaneously, and begin at a fixed time point in larval development, irrespective of larval size (Alberch, et al. 1985; Rose 1999, 2003; Beachy, et al. 2017). Organisms undergo a more extensive remodeling of the feeding apparatus and consequently transform with reduced feeding performance (Deban and Marks 2002). This type of meta-morphosis has evolved approximately three times within salamanders (Bonett, et al. 2014). Abrupt metamorphosers experience the metamorphic vulnerabilities of decreased performance and physiological tolerance, energetic demand, and developmental error due to increased complexity and are thus predicted to have the most extreme levels of constraint on genome expansion associated with metamorphosis.

### Taxon Sampling, Genome Size, and Phylogeny

We analyzed haploid genome size data for 118 species of salamanders (all of which are diploid), including all ten families and 33 of 68 genera, and representing all four life-history regimes (Supplemental Table 1) (AmphibiaWeb 2022). We excluded miniaturized taxa, as miniaturization is associated with decreased genome size independent from any selection imposed by metamorphosis (Decena-Segarra, et al. 2020). Several lineages are facultative paedomorphs; we coded these taxa as metamorphic, as they retain the need to successfully (albeit occasionally) metamorphose (Bonett, et al. 2020).

The dataset includes all species that are represented in the Animal Genome Size Database (www.genomesize.com) and the VertLife database for phylogeny subsampling (www.vertlife.org) (Jetz and Pyron 2018; Gregory 2022). We took the mean genome size where estimates were reported from multiple studies, and natural-log transformed the data to better conform with assumptions of Gaussian errors. We excluded one study that reported consistently higher values than all others (Bachmann 1970). We obtained the 118-species phylogeny by sampling 1,000 ultrametric trees from the pseudo-posterior distribution of the VertLife database (Jetz and Pyron 2018) and computing mean branch lengths using the consensus.edges function in the R package phytools v 0.7-90 (Revell 2012) (Fig. 1). All analyses were conducted in the R statistical computing environment (R Core Team 2020).

**Figure 1.**
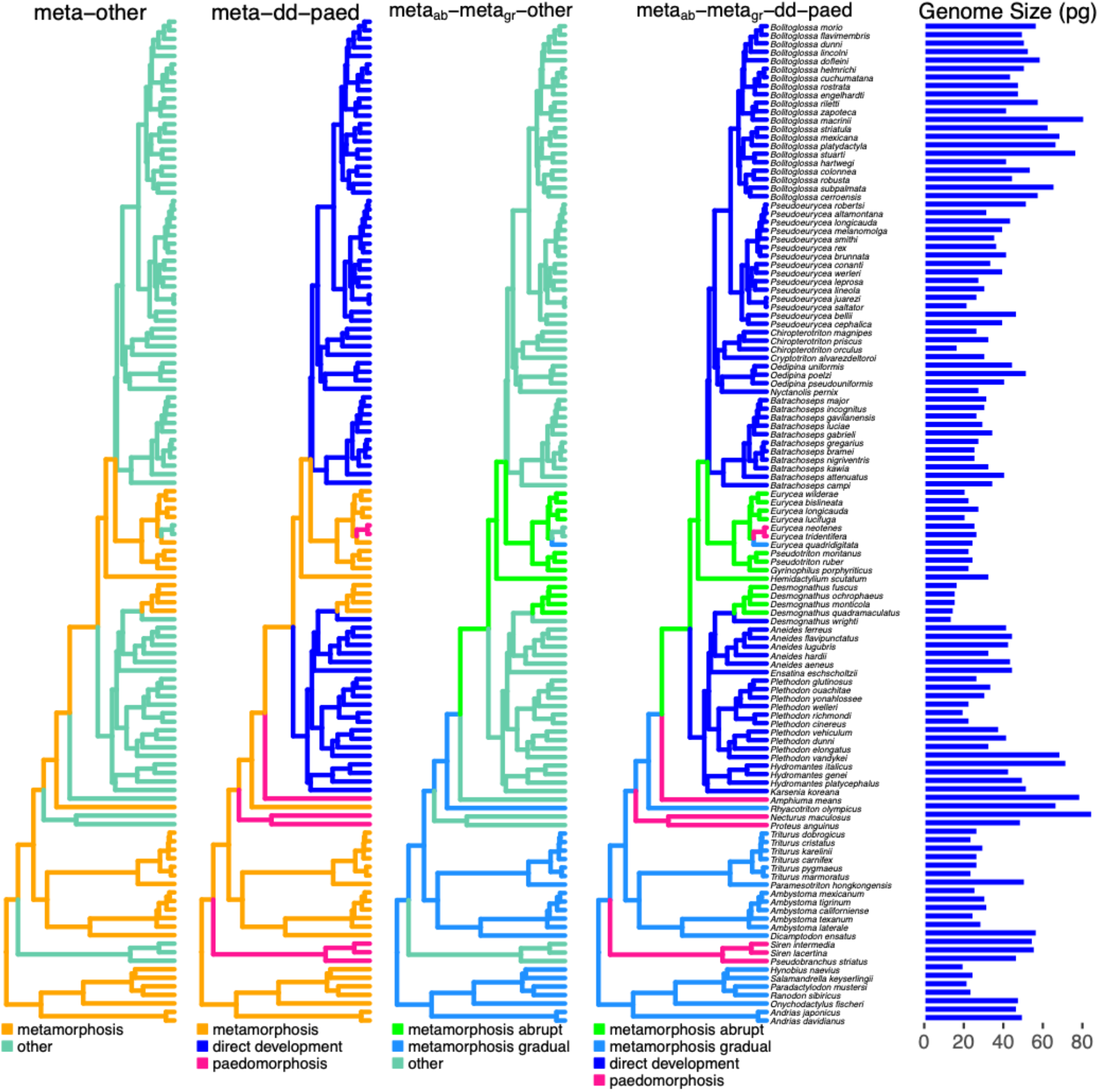
Alternative hypotheses for constraints imposed by development on genome size evolution in salamanders. On each phylogeny (www.vertlife.org), alternative life history regimes are painted in different colors as indicated in each legend (see text). Haploid genome sizes are shown on the right in pg (1 pg = 978 Mb).

### Models of Genome Size Evolution

We modeled genome size evolution using both Brownian motion (BM) and OU models of evolution (Hansen 1997; Butler and King 2004; O’Meara, et al. 2006; Beaulieu, et al. 2012). The BM model is the simplest stochastic model with a single rate parameter σ for *stochastic noise intensity* describing the magnitude of the independent random walks of the trait evolving along the branches of the phylogeny. The multiple-rate BM model allows σ to vary across a phylogeny (O’Meara, et al. 2006).

OU models generalize the BM model by allowing the mean to shift and the variance to narrow. They include a deterministic component of trait evolution that models the tendency to move toward an equilibrium. In contrast to previous work, and to more closely reflect our hypothesis that salamander genome size evolves in response to a balance of mutation pressure and deterministic forces such as selection or constraint, we generalize the notion of “selective optima” to “*deterministic equilibria*”, which encompasses both adaptive and non-adaptive forces affecting the equilibrium (*θ*(*t*)). Similarly, we generalize “selection” to “*deterministic pull”* to encompass non-selective, but directional, evolutionary forces (*α*; e.g., biased mutation pressure and constraint in addition to selection). Mathematically, the model for trait evolution expressed as a differential equation is

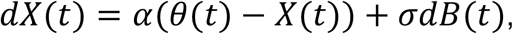

where *θ*(*t*) is the *deterministic equilibrium* for the trait at time *t* and *α* is an evolutionary rate describing the *strength of the deterministic pull* towards that equilibrium. The simplest OU models allow equilibria to vary across the tree, reflecting the evolution of differences in mean phenotype across regimes (Hansen 1997; Butler and King 2004). Further model extensions also allow the strength of the deterministic pull and the stochastic noise intensity to vary across regimes (Beaulieu, et al. 2012).

We formalized five hypotheses for the influence of metamorphosis on genome size evolution: (1) *random evolution*: With no influence of metamorphosis, stochastic evolutionary processes may be sufficient to explain genome size evolution, as represented by BM. The remaining four hypotheses propose different groupings of constraint fit with OU models. (2) *metamorphosis-other*: Metamorphosis imposes a constraint on genome expansion distinct from all remaining life histories, grouped as “other”. (3) *meta-paed-dd*: This hypothesis refines (2) by dividing the “other” category into direct developers and paedomorphs, allowing each to impose distinct constraints on genome size evolution. (4) *meta_abrupt_-meta_gradual_-other*: Alternatively, we may refine (2) by keeping the “other” category of non-metamorphosers, but differentiating the two groups of metamorphosers, recognizing that abrupt (*meta_abrupt_*) and gradual (*meta_gradual_*) metamorphosis each impose distinct constraints on genome size evolution. (5) *meta_abript_-meta_gradual_-paed-dd*: Each category imposes distinct constraints on genome expansion (Fig. 1).

We tested 21 models that varied in the number of parameters used to explore these five biologically-inspired hypotheses. The simplest model allows the equilibria to vary with life history regime while modeling a single stochastic noise intensity (σ) and deterministic pull strength (*α*) across all species. Additional sub-hypotheses fit evolutionary models with separate stochastic noise intensities (*σ*_*i*_) for each regime (Table 1). More complex models with multiple deterministic pull strengths (*α*_*i*_) produced fitting errors or nonsensical parameter estimates or likelihoods, so we do not consider them further (see Supplementary Information; Table 2). We fit the remaining 13 models with the ancestral plethodontid node assigned to be either metamorphosing or direct-developing (Bonett, et al. 2014) and found that the choice made no qualitative difference (Supplementary Information). Model fitting and parameter estimation were carried out using OUwie (Beaulieu, et al. 2012).

**Table 1.**
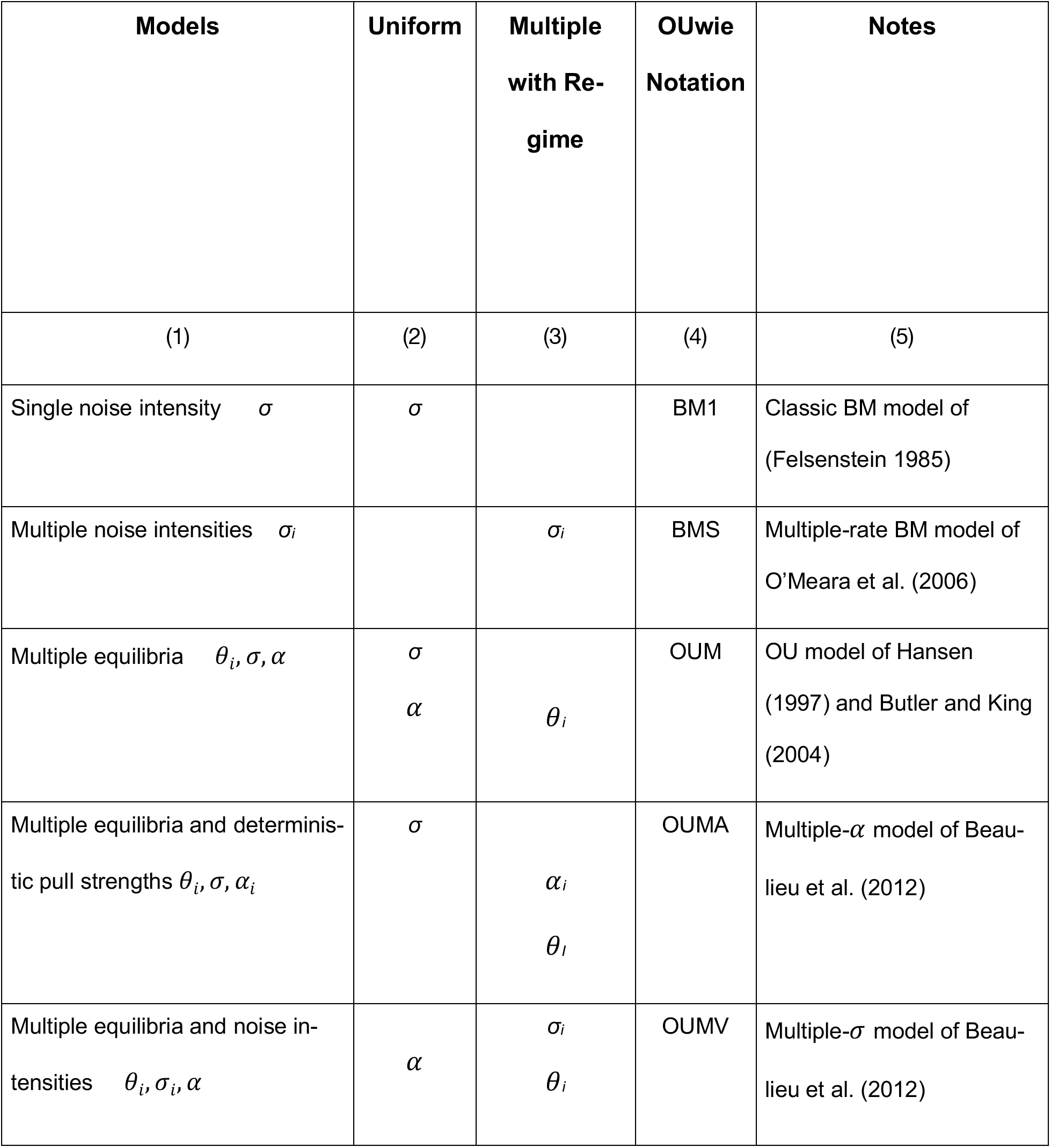
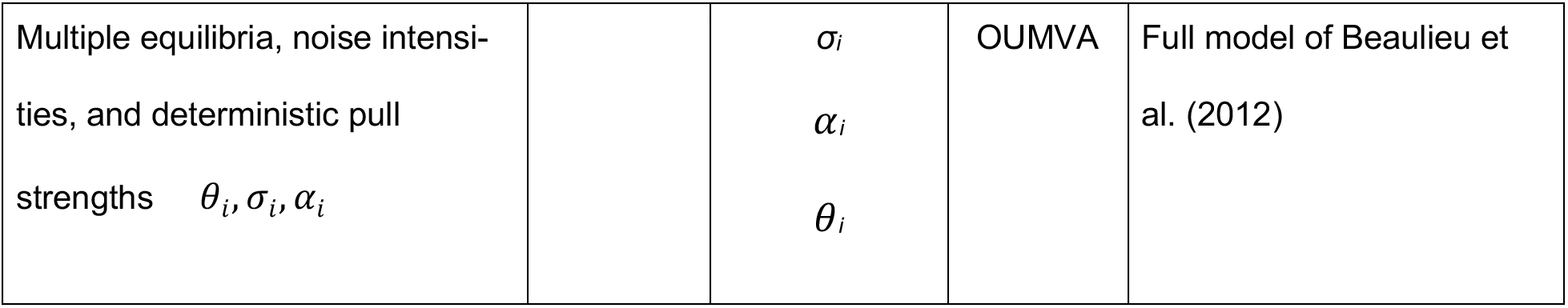
BM and OU models with single or multiple parameters used to fit the data. Numbers in parentheses specify (1) model parameters and notation, (2) parameters that remain constant across the phylogeny, (3) parameters that vary with shifts in life history regime, (4) OUwie model notation, and (5) notes for the model implementations and citations.

### Model Comparison

We compared the fit of each of the models using the Akaike Information Criterion corrected for small sample size (AIC_c_). Because AIC_c_ differences can favor more complex models even when a simpler one is correct, we performed model selection bootstrap analysis (phylogenetic Monte Carlo; (Boettiger, et al. 2012). We additionally evaluated the support for hypotheses in six pairwise comparisons (Fig. 2) which assess and progressively refine the strength of evidence for successive levels of increased model complexity as well as the power to detect differences in model support. Support for the more complex model in each pair implies:

A. *BM vs. metamorphosis-other*: metamorphosis imposes a constraint on genome expansion;
B. *meta-other vs. meta-dd-paed*: distinct constraints are imposed by the different non-metamorphosing strategies, direct development and paedomorphosis;
C. *meta-other vs. abrupt-gradual-other*: abrupt metamorphosis imposes a distinct constraint from gradual metamorphosis;
D. *meta-dd-paed vs. abrupt-gradual-dd-paed*: abrupt metamorphosis imposes a distinct constraint from gradual metamorphosis, after accounting for differences in non-metamorphosing strategies;
E. *abrupt-gradual-other vs. abrupt-gradual-dd-paed*: non-metamorphosing strategies impose unique constraints, after accounting for differences between abrupt and gradual metamorphosis;
F. after identifying *meta_abrupt_-meta_gradual_-paed-dd* as the best-fitting hypothesis, we further explored the power to discriminate between multiple versus single stochastic noise intensity parameters.

**Figure 2.**
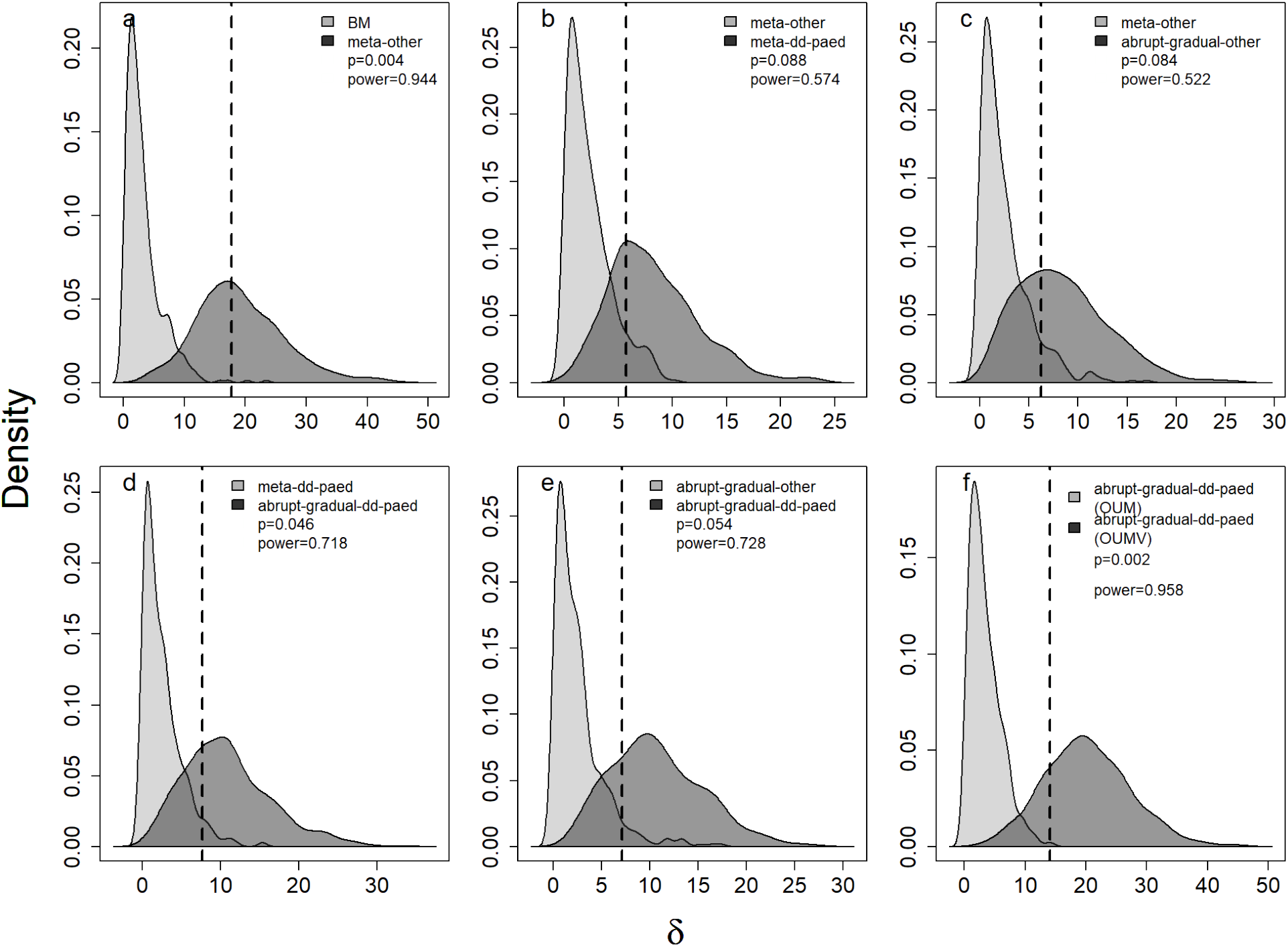
The power to discriminate among competing life-history regime models. Each panel evaluates the support for a different, more complex, hypothesis relative to a simpler one (panel keys). Pairwise bootstrap distributions of the likelihood difference (*δ*) calculated by generating 500 datasets under each of two competing life history regime models at their MLE parameter estimates, refitting the two models, and computing *δ*. All comparisons are between multiple equilibria, multiple noise intensity models. Assuming the simpler model is the truth, the probability density of *δ* is in light gray, while the density of the likelihood difference assuming the complex model is in dark gray. The dashed line gives the observed value (*δ*_obs_) from fitting the actual genome size data. The reported *p-*value is the fraction of the light gray distribution that lies to the right of *δ*_obs_; the power is the fraction of the dark gray distribution that lies to the right of the 95^th^ percentile of the light gray distribution.

For each comparison, we computed the observed likelihood difference,

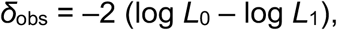

where *L*_0_ is the likelihood of the simpler model and *L*_1_ is the likelihood of the more complex model. We used these parameters and stochastic simulation to compute approximate *p-*values and power.

Determining whether *δ*_obs_ is significantly different from a null expectation requires an approximate *p-*value — the probability of observing *δ*_obs_ *if the simpler model were true*. That is, we need to compare the value *δ*_obs_ to the distribution of *δ* values under the simpler model. To create this distribution, we generated 500 datasets by simulating the simpler model at its MLE parameter estimates; we then fit both the simpler and more complex models to each simulated dataset and computed the values of *δ*, producing a null distribution of *δ* assuming the simpler model. We compared the observed value of *δ* to this null distribution to calculate an approximate *p*-value.

Power conveys the (desirable) probability of rejecting the simpler model when the more complex model is true. To estimate power, we generated 500 datasets by simulating the more complex model at its MLE parameter estimates; we then fit the two models and computed the values of *δ*. The fraction of these *δ* values that are greater than the 95% quantile of the distribution generated under the simpler model (described above) gives an estimate of power. All data and code necessary to carry out the analysis in this manuscript can be found at https://github.com/claycressler/genomesize and in Supplemental Material.

## Results

The best-fitting model for salamander genome size evolution accounted for four regimes: both abrupt and gradual metamorphosis, paedomorphosis, and direct development (*meta_abrupt_-meta_gradual_-paed-dd*; Table 2) under an OU model that allowed both equilibrium genome size and noise intensity to vary across these regimes (θ_i_, σ_i_, α, Table 2). The three-regime *meta_abrupt_ -meta_gradual_ -other* (θ_i_, σ_i_, α) and *meta-dd-paed* (θ_i_, σ_i_,α) hypotheses also had marginal support (*Δ*AIC_c_ ≤ 3).

Several authors have recommended parametric and model selection bootstraps to assess OU models, as model selection tends to be robust while parameters can be difficult to estimate accurately (Boettiger, et al. 2012; Ho and Ané 2013; Ho and Ané 2014; Cressler, et al. 2015). We found no evidence of identifiability issues; AIC_c_ values correspond with parametric bootstrap results, and parameter estimates and confidence intervals are well-behaved (Table 3, Fig. 2,3). We furthermore avoid interpreting point estimates of parameters, limiting our discussion to features of the best models, relative rankings of parameter estimates across regimes bolstered by AIC_c_-derived confidence intervals, and comparison of variation across models.

**Table 2.** Model comparison statistics. Best model (interrogated by bootstrap, Fig. 2) indicated in bold. Model parameterizations are indicated by: σ = Brownian motion; σ_*i*_ = Brownian motion with multiple noise intensities; θ_*i*_, σ, α = OU model with multiple equilibria; θ_*i*_, σ_*i*_, α = OU model with multiple equilibria and multiple noise intensities.

| | $\Delta AIC_c$ | | | |
| --- | --- | --- | --- | --- |
|  | OU Models |  | BM Models |  |
| Hypotheses | $\theta_i, \sigma_i, \alpha$ | $\theta_i, \sigma, \alpha$ | $\sigma_i$ | $\sigma$ |
| *meta*<sub>abrupt</sub> - *meta*<sub>gradual</sub> - *paed* - *dd* | **0** | 7.1 | 7.5 |  |
| *meta*<sub>abrupt</sub> - *meta*<sub>gradual</sub> - *other* | 2.5 | 6.9 | 9.5 |  |
| *meta* - *paed* - *dd* | 3.0 | 8.8 | 5.3 |  |
| *metamorphosis* - *other* | 4.2 | 7.5 | 7.6 |  |
| Brownian motion |  |  |  | 15.5 |

**Table 3:**
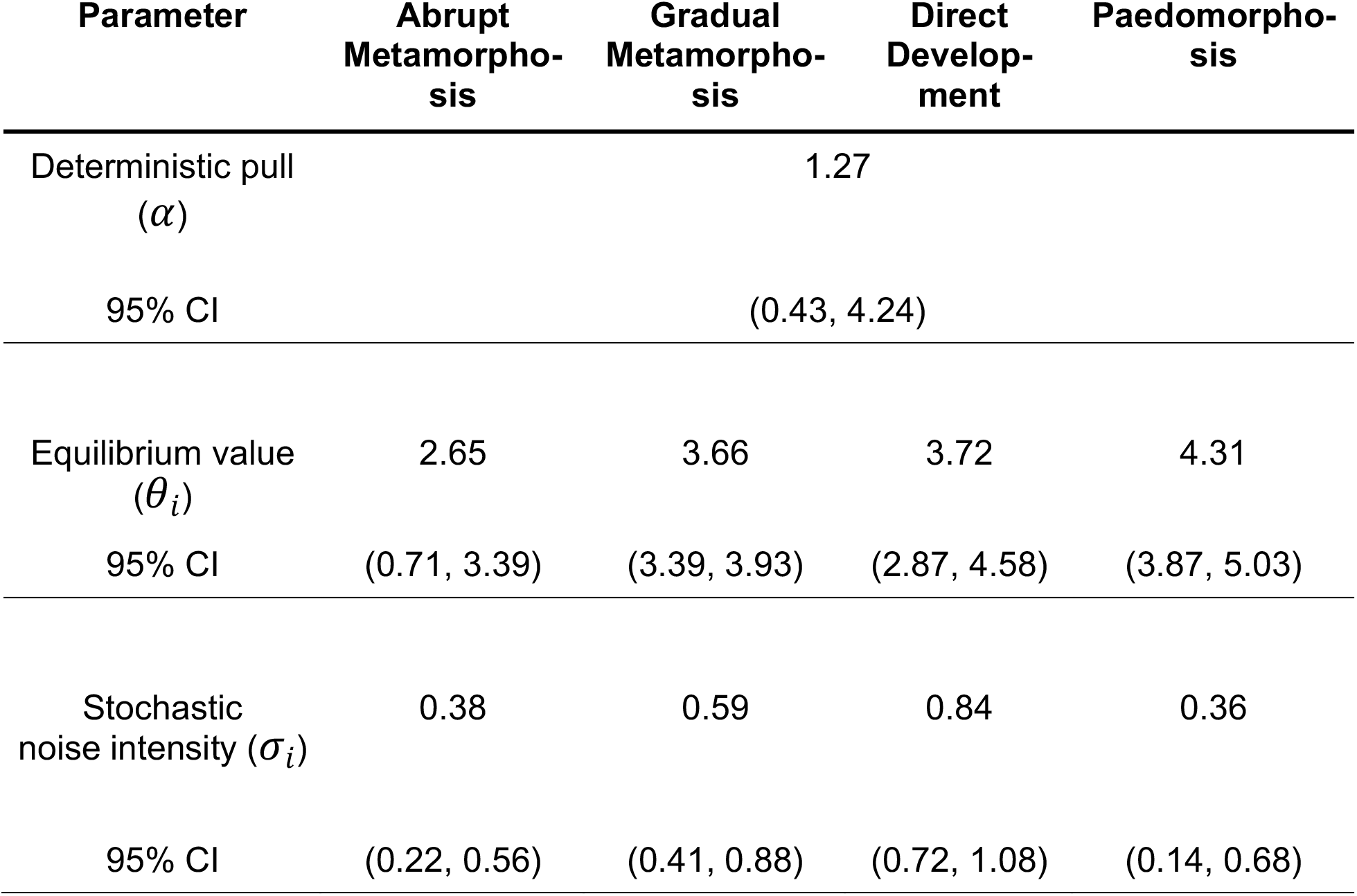
Maximum likelihood parameter estimates and parametric bootstrap confidence intervals for the best-fitting model (*meta_abrupt_-meta_gradual_-dd-paed θi*, *σi*, *α*: separate equilibrium values and noise intensities for lineages in the four life history regimes: dd = direct development, meta_*abrupt*_ = abrupt metamorphosis, meta_*gradual*_ = gradual metamorphosis, paed = paedomorphosis)

**Figure 3.**
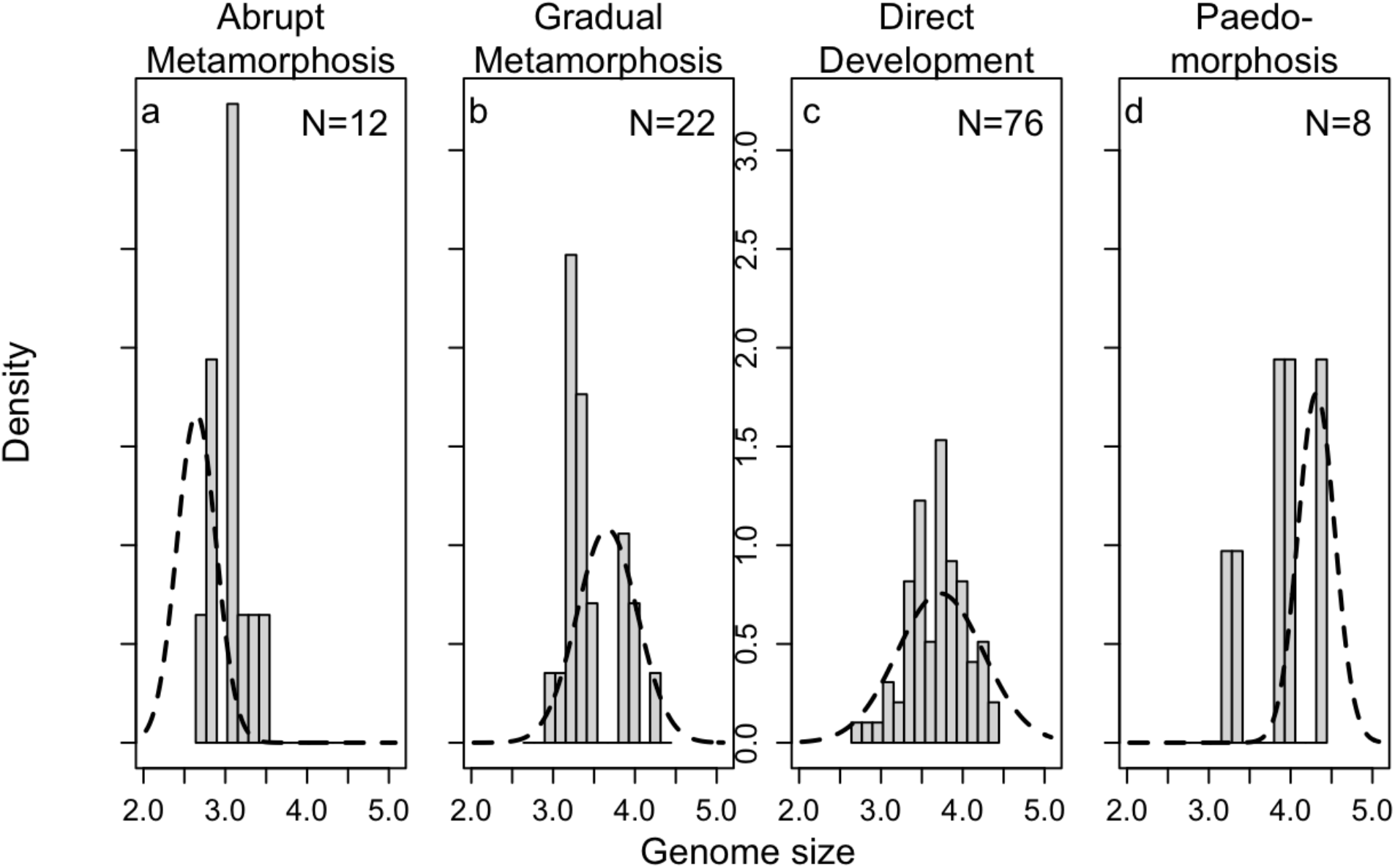
Comparison of the observed distribution of genome sizes for salamanders in each regime (bars) against the stationary distribution predicted by the best-fitting parameter set (dotted lines, Table 3). The stationary distribution is normal with mean *θ* and variance *σ*^2^/(2*α*), where the values of *θ* and *σ* vary across regimes. The x-axis is naturallog transformed genome size.

We reject a purely stochastic hypothesis for genome size evolution based on both the results of model fitting and interrogation by parametric bootstrap (Table 2, Fig. 2). The BM model had the highest AIC_c_, and a model that accounts for metamorphosisvs. others with separate equilibrium and noise intensity values was far superior to a purely neutral model (Table 2; Fig. 2a). Models that separate non-metamorphosing strategies into direct development and paedomorphosis (comparing *metamorphosis-other* to *meta-dd-paed*; Fig. 2b) or separate metamorphosing strategies into gradual and abrupt metamorphosis (comparing *metamorphosis-other* to *meta_abrupt_-meta_gradual_-other*; Fig. 2c) had only moderate power to reject the simpler *metamorphosis-other* hypothesis. However, models that included all four selective regimes (*meta_abrupt_-meta_gradual_-dd-paed*) had high power to reject any three-regime hypothesis (Fig. 2d,e). These model selection bootstrapping results support our conclusion, based on the AIC_c_ values obtained from fitting the real data, that these life history regimes have different deterministic equilibria (Table 3), and moreover, allowing noise intensity to vary among regimes is strongly supported (Fig. 2f). Therefore, we have compelling evidence that observed differences in genome size between abrupt metamorphosers, gradual metamorphosers, direct developers, and paedomorphs (Fig. 3) reflect differences in the balance of evolutionary forces shaping genome size in each regime.

Parameter estimates for the best-fitting model are presented in Table 3. Abrupt metamorphosers have the smallest genome size equilibrium, gradual metamorphosers and direct developing salamanders are intermediate, and paedomorphic salamanders have the largest (Table 3), with multiple categories of stochastic noise intensity. To help visualize what this best-fitting model tells us about genome size evolution, we compared the distribution of observed genome sizes for species in each selective regime to the expected stationary distribution, given the model estimates of *θ*, *α*, and *σ* for each regime (Fig. 3). In particular, (Ho and Ané 2013) showed that the stationary distribution of an OU process will be a normal distribution with mean *θ* and variance *σ*^2^/(2*α*), providing a useful way to visualize the model fit that captures aspects of all parameters. In the direct-developing regime (which is the most common across the tree), the expected distribution completely overlaps the observed (Fig. 3c). The observed and expected distributions of genome size for gradual metamorphosers are broadly overlapping, although the mean of the expected distribution is slightly higher than the observed (Fig. 3b); similar patterns are observed for paedomorphs (Fig. 3d). In contrast, the expected mean is smaller than almost all abrupt metamorphosing salamander genome sizes (Fig. 3a). Furthermore, this best-fit model predicts that the variance of the trait distribution for each regime is well-predicted by *σ*^2^/(2*α*), which varies by regime because of differences in the noise parameter *σ_i_*. Direct developers have larger stochastic noise in genome size than abrupt metamorphosers or paedomorphs, with gradual metamorphosers intermediate (Table 3, Fig. 3).

## Discussion

### Genome Size Evolution Across Different Life Histories

Genome size differs among metamorphic life history strategies in salamanders. Salamanders that undergo abrupt metamorphosis are experiencing deterministic pull downward, suggesting that abrupt metamorphosis is driving the evolution of smaller genome size (smaller equilibrium value than observed, Figure 3a). Gradual metamorphosers and direct developers reach larger genome sizes than abrupt metamorphosers, but we see little evidence from the best-fit model of any tendency towards either genome size contraction or expansion (Fig. 3b,c). When metamorphic remodeling is largely removed from life history, genomes are expected to be unconstrained and permissive to TE accumulation, consistent with our finding that paedomorphic lineages are experiencing deterministic pull toward genome expansion (Fig. 3d). Although metamorphosis has been suggested to influence the evolution of genome size, this hypothesis has rarely been demonstrated in a broadly comparative manner, and the relevant features of metamorphosing organisms that exert evolutionary pressure on genome size have been under-studied (but see Gregory 2002b; Bonett, et al. 2020). We discuss how three classes of metamorphic vulnerabilities or their interaction may constrain biased TE accumulation and, thus, shape genome size diversity across life history regimes in salamanders: (1) performance and physiological handicap, (2) energetic limitation, and (3) developmental error during metamorphosis. We track the influence of these evolutionary pressures in qualitative form in Table 4.

**Table 4.** Mutation pressure and metamorphic vulnerabilities shaping genome size in different life history regimes

| Evolutionary Pressure | Effect on Genome Size | Abrupt Metamor-phosers | Gradual Metamor-phosers | Direct Developers | Paedomorphs |
| --- | --- | --- | --- | --- | --- |
| TE accumulation bias | ↑ | ✓<sup>a</sup> | ✓ | ✓ | ✓ |
| Performance or Physiological Handicap | ↓ | ✓ | ✓ |  |  |
| Energetic Limitation | ↓ | ✓✓ |  | ✓ |  |
| Developmental Error | ↓ | ✓✓ | ✓ | ✓ |  |

<sup>a</sup>A single checkmark vs. two checkmarks indicates a difference in the intensity of a metamorphic vulnerability across life history regimes.

### Performance and Physiological Handicap

Performance handicaps are expected based on the classic and widely-cited work on frogs; metamorphosing individuals can neither hop nor swim effectively and are subject to increased predation (Wassersug and Sperry 1977; Arnold and Wassersug 1978). In contrast, metamorphosing salamanders do not suffer locomotor handicaps (Landberg and Azizi 2010). However, some metamorphosing salamanders (*Gyrinophilus porphyriticus*) do suffer a performance handicap in accessing shelter during critical environmental fluctuations; their decreased ability to exploit stream habitat refugia during extremely high or low flow rates increases mortality (Lowe, et al. 2019). In addition, several species of salamanders have decreased critical thermal maxima during metamorphosis (*Ambystoma tigrinum*, *Notophthalmus viridescens*) (Hutchison 1961; Delson and Whitford 1973). Thus, decreased performance and physiological tolerance likely contribute to the increased mortality experienced by salamanders during metamorphosis (Peterson, et al. 1991), selecting for rapid metamorphosis and constraining genomes and cells to smaller sizes. However, these vulnerabilities apply to all metamorphosers, accounting for their smaller genome sizes relative to direct developers and paedomorphs, but not for the differences between abrupt and gradual metamorphosers (Table 4).

### Energetic Limitation

Vulnerability of metamorphosing individuals to energetic limitation is important despite low overall metabolic rates across salamanders (Gatten, et al. 1992), and furthermore affects the metamorphic strategies differently. The most strongly affected are abrupt, synchronous metamorphosers. There is evidence that these larvae are food-limited in nature; individuals grow more slowly in the field than those fed *ad libitum* in the lab (Beachy 1995). In addition, the timing of the onset of abrupt metamorphosis is fixed, irrespective of food availability, larval growth rate, or body size (Beachy, et al. 2017; Beachy 2018). Thus, these animals lack the flexibility to delay metamorphosis until they reach an optimal size — unlike metamorphosing frogs, arthropods, and salamanders that undergo gradual metamorphosis — resulting in the possibility that transformation is forced at a smaller body size with lower energy reserves, yielding reduced fitness (Wilbur and Collins 1973; Beachy, et al. 2017). Once abrupt metamorphosis begins, feeding ceases, although there are mixed anecdotal reports of transforming organisms feeding or attempting to feed in the lab (Deban and Marks 2002). Metamorphosis itself does not increase energetic demand, as evidenced by gradually metamorphosing salamanders, in which metabolic rate remains unchanged during the transformation (Vladimirova, et al. 2012). However, the transformation is more extensive in abrupt metamorphosing salamanders, and frogs experience elevated metabolic rates during their extensive meta-morphosis (Orlofske and Hopkins 2009; Wright, et al. 2011). Thus, abrupt metamor-phosers are at risk of initiating a costly sequence of developmental events with sub-optimal energy reserves and impaired capacity to feed (Deban and Marks 2002; Vladimirova, et al. 2012; Beachy, et al. 2017). In contrast, gradual metamorphosers are not affected energetically, as they can feed throughout metamorphosis as well as delay its onset. In direct-developing lineages, some or all of the developmental steps of meta-morphic remodeling occur inside the egg at the end of embryogenesis, in sequences that deviate from those of metamorphosers to varying degrees (Alberch 1989; Wake and Hanken 1996; Marks 2000; Rose 2003; Kerney, et al. 2012). The energy to fuel metamorphic repatterning comes from yolk stores which, although relatively large in direct developers, are still finite (Wake and Hanken 1996; Gregory 2002b), suggesting the potential for some energetic vulnerability. However, direct developers do not have smaller genome size equilibria than gradual metamorphosers, suggesting that any constraint imposed by energetic limitation in direct developers is weak. Abrupt metamor-phosers have the smallest genome size equilibrium, consistent with energetic limitation exerting the strongest selection for rapid metamorphosis and therefore the strongest constraint on genome size (Table 4).

### Developmental Error

Abrupt and gradual metamorphosers differ in both metamorphic synchronicity and extent of transformation, leading to differential vulnerability to developmental errors. Abrupt metamorphosis entails the synchronous execution of a large number of developmental events, e.g.: the chondrification and ossification of the nasal capsule, dermal bones of the skull, upper jaw, and palate; remodeling of the skin; and gill resorption. The internal organs are likely also abruptly remodeled because they are under the same hormonal control (Rose 1996; Rose 1999, 2003). Some systems undergo a more radical transformation altogether; larval elements undergo cell death, and adult structures form *de novo* rather than being remodeled from larval elements. For example, the hyobranchial apparatus (i.e. tongue skeleton) in abrupt metamorphosers transforms this way, enabling the formation of projectile and, in some species, ballistic tongues (Alberch, et al. 1985; Alberch and Gale 1986). These transformations require precise spatial coordination of the cell-autonomous programs and cell-cell interactions that drive cell death, growth, and differentiation, often in very close proximity (Alberch, et al. 1985; Schreiber, et al. 2009; Ishizuya-Oka 2011). Temporal organization is also critical; induction events (e.g. epithelial-mesenchymal interactions) that occur out of sequence result in developmental anomalies (Hanken and Hall 1993). The evolution of abrupt metamorphosis increased the complexity of spatiotemporal coordination of cell-cell interactions relative to gradual metamorphosers. We found that abrupt metamorphosers have distinctly smaller genome sizes, consistent with more intense vulnerability to developmental error (Table 4). This result is broadly consistent with a similar pattern in insects, where holometabolous lineages (i.e. those that undergo a complete metamorphosis) have smaller genome sizes than hemimetabolous lineages (i.e. those that undergo incomplete metamorphosis), which has been interpreted as evidence for stronger constraint on genome size with greater developmental complexity (Gregory 2002b; Alfsnes, et al. 2017).

Direct developers undergo repatterning at a developmental stage involving much less tissue and fewer cells, therefore requiring less spatiotemporal cell-cell coordination. Although this simplifies the overall developmental process, it also introduces the potential for error because stochastic noise in developmental processes (e.g. cell migration) can have larger phenotypic effects when each cell represents a larger proportion of an incipient structure. One of the best illustrations of the mechanistic consequences of low cell numbers on development is the salamander genus *Thorius*, characterized by large genomes, small bodies, and thus low cell numbers. Skeletal and cartilaginous elements in the limbs and skull form embryonically from precartilage condensations, which are tight aggregates of mesenchymal cells. Up to 70% of *Thorius* individuals show left-right asymmetry in the arrangement of carpal or tarsal elements in the limbs and/or anterior elements in the skull, and similar variation exists among individuals (Hanken 1982; Hanken 1984). This degree of variability demonstrates that the outcome of precartilage condensation is subject to stochastic noise in the cellular processes involved in cell aggregation (i.e. cell movement, cell-cell adhesion, cell-extracellular matrix interaction) (Chatterjee, et al. 2020; Glimm, et al. 2020).

More generally, vulnerability to stochastic noise is expected to impose stronger constraints on genome expansion when developmental sequences are more complex. Limited data from direct developers support this hypothesis. For example, in the genus *Bolitoglossa*, the formation of larval hyobranchial apparatus components has been lost from ontogeny, leading to a simpler developmental sequence in which adult structures are formed without larval precursors (Alberch 1989; Wake and Hanken 1996). Other taxa (e.g. *Desmognathus* and the eastern *Plethodon*) retain a more complex developmental sequence with more of the larval and metamorphorphic stages of hyobranchial development (Wake and Hanken 1996; Marks 2000; Kerney, et al. 2012); as predicted, they have smaller genome sizes (~15 Gb and ~25 Gb for *Desmognathus* and the eastern *Plethodon*, respectively, versus ~50 Gb for *Bolitoglossa*).

Both direct developers and abrupt metamorphosers experience risks for developmental error, but direct developers have a larger genome size equilibrium (Table 3), supporting weaker developmental error constraints (Table 4). In addition, direct developers have the largest expected variance in genome size, which presents a new hypothesis — that diverse developmental sequences within this life history regime also differ in vulnerability to error and, thus, constraint on genome expansion.

### Paedomorphosis and the Loss of Metamorphic Constraints

Despite their freedom from metamorphosis-induced constraints, paedomorphs may still face other evolutionary pressures that prevent their genomes from reaching ever-larger sizes. The decreased surface-area-to-volume ratio that accompanies increased cell size imposes a functional limit that salamanders may well have reached (Chan and Marshall 2010), as their cells are among the largest in animals (Horner and Macgregor 1983). In addition, the ability of adult salamanders to regenerate limbs and organs declines dramatically in paedomorphs with the largest genomes, suggesting another fitness consequence of extreme genome expansion (Scadding 1977; Sessions and Wake 2021).

In the past, huge cells have been proposed as adaptive because salamanders and lungfishes have the lowest metabolic rates and the largest genomes/cells among vertebrates. This correlation led to the “frugal metabolic strategy” hypothesis (Szarski 1983; Olmo, et al. 1989), under which paedomorphs would show the greatest degree of adaptation for this trait. However, more recent studies failed to find a clear relationship between genome or cell size and metabolic rate, both within salamanders (Licht and Lowcock 1991) and more generally (Uyeda, et al. 2017; Gardner, et al. 2020). Rather than adaptation towards larger genome size, our results support the idea that loss of metamorphosis has released constraints against genome expansion in paedomorphs.

### Summary of Metamorphic Vulnerabilities and Genome Size Evolution

Abrupt metamorphosers are challenged with every metamorphic vulnerability: decreased performance and physiological tolerance, energetic limitation, and developmental error, providing the explanation for their smallest genome sizes (Table 4). In contrast, paedomorphs are free from all of these vulnerabilities and have the largest genome sizes (Table 4). Direct developers have a similar genome size equilibrium to gradual metamorphosers, but with larger variation (Table 3), which we propose reflects diversification of direct developmental pathways. Gradual metamorphosers avoid developmental error at the cost of decreased performance and physiological tolerance whereas direct developers avoid performance handicap at the cost of some susceptibility to stochastic noise in development. More generally, these results illustrate how genome size may be shaped by different constraints across taxa that are important to assess in order to understand the major drivers of genome size evolution across the tree of life (Knight, et al. 2005; Carta and Peruzzi 2016; Alfsnes, et al. 2017; Roddy, et al. 2019).

### OU Models Evaluate the Balance of Evolutionary Forces

While we have no model tailored to biased evolutionary increase opposed by constraining forces, the OU class of models (of which BM is a subset) is a powerful tool that can differentiate among biologically-motivated, nuanced hypotheses. For example, because TE accumulation is stochastic, one might expect the BM model to be a good candidate for purely stochastic genome size evolution. However, this is not the case for salamanders because there is bias for TE gains over losses, rendering BM a poor fit. While it is widely recognized that OU models provide deterministic components that allow modeling evolutionary shifts in the means of phenotypes, it is less appreciated that they also provide a way to model the information content in the variances. The difference between a purely stochastic (BM) model and one that has any degree of deterministic pull (OU) is that the variance of a BM model will grow unbounded over time, whereas the variance in a model with deterministic pull will not (Hansen and Martins 1996; Butler and King 2004). Along these lines, our finding that allowing both stochastic noise intensities and equilibria to vary across regimes produces a model with substantially better fit than either set of parameters alone is very biologically informative. While we have three broad categories of genome size equilibria, with abrupt metamorphosers distinctly smaller and paedomorphs larger (Table 3), the metamorphic categories also differ substantially in stochastic noise intensity. The smaller equilibrium value and noise intensity of abrupt metamorphosers suggests that this regime is highly constrained with deterministic pull downward, even in comparison to gradual metamorphosers (Table 3, Fig. 3a vs. b). Direct developers have the highest estimated noise intensity, suggesting that the evolution of diverse developmental sequences resulted in a range of vulnerabilities to error and, thus, variable constraints on genome expansion (compare confidence intervals in Table 3, Fig. 3c vs. d). Thus, both the mean and variance of the evolutionary process shed light on the balance of forces acting on genome size across life histories.

In general, OU models are conceptualized as models for stabilizing selection, with the *θ* parameters interpreted as trait optima towards which individual species are evolving. However, this interpretation does not fit well with the biology of genome size. As the variability in genome size is strongly determined by the quantity of nearly-neutral noncoding DNA, the notion of an “optimal” genome size has little meaning (but see Cavalier-Smith 2005). More realistically, there is a range of permissible variation within which species can vary.

We propose that the strong support for OU models in our analysis reflects a balance between the biased stochastic forces driving genome size upward and evolutionary constraint acting to limit genome size. That is, rather than interpreting Fig. 3b as evidence that gradual metamorphosing salamanders have genomes near their “optimal”sizes, we interpret it as showing that these salamanders have settled on a genome size distribution that is balanced between stochastic TE dynamics tending to bias genome size upward and selection on metamorphosis imposing evolutionary constraint against further size increase. Paedomorphs have the largest genome size equilibria of all, and our model indicates that they are evolving with a deterministic pull toward even larger size. There is no compelling adaptive interpretation for this genome size “optimum” larger than all other vertebrates, although some have tried (see above); rather, we interpret it as a balance between upwardly-biased TE accumulation and functional constraints not considered here (e.g. an upper limit on cell size). This is a novel way of interpreting the results of comparative analysis, but one that is supported by our understanding of the biology of the system. Our results thus suggest that OU models can potentially be used to detect other evolutionary processes beyond adaptation towards an optimum, which broadens their applicability to the study of traits that do not evolve in response to strong selection by a single factor.

